# Hypoxic environment promotes barrier formation in human intestinal epithelial cells through regulation of miRNA-320a expression

**DOI:** 10.1101/483206

**Authors:** Stephanie Muenchau, Rosalie Deutsch, Thomas Hielscher, Nora Heber, Beate Niesler, Megan L. Stanifer, Steeve Boulant

**Affiliations:** Schaller research group at CellNetworks, Department of Infectious Diseases,Virology, Heidelberg University Hospital, Germany; Division of Biostatistics, German Cancer Research Center (DKFZ), Heidelberg, Germany; Department of Human Molecular Genetics, Institute of Human Genetics, University of Heidelberg, Heidelberg, Germany; Research Group "Cellular Polarity of Viral Infection", German Cancer Research Center (DKFZ), Heidelberg, Germany

**Keywords:** barrier function, hypoxia, tight junctions, miRNA, intestinal epithelial cells

## Abstract

Intestinal epithelial cells (IECs) are exposed to the low-oxygen environment present in the lumen of the gut. These hypoxic conditions are on one hand fundamental for the survival of the commensal microbiota, and on the other hand, favor the formation of a selective semipermeable barrier allowing IECs to transport essential nutrients/water while keeping the sterile internal compartments separated from the lumen containing commensals. The hypoxia-inducible factor (HIF) complex, which allows cells to respond and adapt to fluctuations in oxygen levels, has been described as a key regulator in maintaining IEC barrier function by regulating their tight junction integrity. In this study, we sought to better evaluate the mechanisms by which low oxygen conditions impact the barrier function of human IECs. By profiling miRNA expression in IECs under hypoxia, we identified miRNA-320a as a novel barrier formation regulator. Using pharmacological inhibitors and short hairpin RNA-mediated silencing we could demonstrate that expression of this miRNA was HIF-dependent. Importantly, using over-expression and knock-down approaches of miRNA-320a we could confirm its direct role in the regulation of barrier functions in human IECs. These results reveal an important link between miRNA expression and barrier integrity, providing a novel insight into mechanisms of hypoxia-driven epithelial homeostasis.

## Introduction

The human gastrointestinal (GI) tract is the organ forming the largest barrier towards the external environment and a key player in nutrient absorption (1). It is made of a monolayer of epithelial cells separating the *lamina propria* from the lumen of the gut. This epithelium on the one hand allows for the translocation of nutrients, water and electrolytes from the lumen to the underlying tissue and, on the other hand, builds up a tight barrier to prevent penetration of commensal bacteria and potential harmful microorganisms (bacterial and viral) to the *lamina propria* (1). Although these luminal microorganisms have well characterized beneficial functions for the host, they can represent a risk when epithelial barrier and gut homeostasis are disrupted. Altered barrier functions increase the risk of enteric pathogen infection and can lead to the dysregulation of the mechanisms leading to the tolerance of the commensals which ultimately can lead to inflammation of the GI tract and the development of chronic diseases like inflammatory bowel disease (IBD), including Crohn’s disease (CD) and ulcerative colitis (UC) (2, 3). Multiple cellular strategies are utilized to physically separate the content of the gut lumen from the host. First, goblet cells and Paneth cells in the mucosal lining secrete mucus together with antimicrobial and antiviral peptides which forms a layer of separation between the intestinal epithelial cells and the luminal content of the digestive tract (4–6). Second, epithelial cells polarize and express tightly juxtaposed adhesive junctional complexes between neighbouring cells. These junctional complexes are composed of integral transmembrane proteins that are linked via intracellular scaffoldings proteins to the actin cytoskeleton (7). This tight organization of intestinal epithelial cells (IECs) inhibits paracellular diffusion of ions and other solutes as well as antigenic material (8). The junctional complex therefore is essential for establishing and maintaining the barrier function of the mucosal layer and is composed of tight and adherens junction proteins such as claudins, occludin, junctional adhesion molecule-A (JAM-A), tricellulin, zona occludens-1 (ZO-1) and E-cadherin (8). The interaction between the different tight junction and adherens junction proteins thus creates a tight epithelial barrier and determines selective permeability through the intestinal epithelium.

Within the physiological organization of the GI tract, an important but often overlooked parameter is the low oxygen level present in the lumen of the gut. This environment is fundamental for the survival of many commensals. Within the complex 3D organization of the crypt-villus axis, the tip of the villi protrudes into the low oxygen (1-2%) environment of the gut (hypoxic environment) (9). Conversely, within of the mucosal lining, oxygen-rich blood vessels are located in the subepithelium, providing the stem cell containing crypts with a high oxygen content of around 8-21% (normoxic environment) (10, 11). Besides this oxygen gradient among the intestinal epithelium, the subepithelium of the GI tract is also exposed to daily fluctuations in oxygen content. After food ingestion, the intestinal blood flow increases and the oxygen content in the subepithelium rises up to 40-64%, but can also decrease below 8% under fasting conditions (12, 13).

Cells respond to the hypoxic environment by specifically regulating the expression of hundreds of genes through the major hypoxic-induced transcription factor hypoxia inducible factor (HIF) (14). HIFs are heterodimeric transcription factors that are composed of a constitutively expressed HIF-β subunit and one of the three oxygen-regulated alpha subunits (HIF-1α, HIF-2α or HIF-3α) (15). Under normoxic conditions, HIF-1α is rapidly hydroxylated at specific proline residues by different prolyl hydroxylases (PHD’s), leading to binding to the E3 ubiquitin ligase containing the von Hippel-Lindau (VHL) tumor suppressor protein, polyubiquitination and subsequent proteasomal degradation of the protein (16). Under hypoxic conditions, lack of substrates such as Fe^2+^, 2-oxoglutarate and O_2_ inhibits hydroxylation (17), therefore stabilizing HIF-1α and leading to dimerization with its constitutively expressed β -subunit (HIF-1β), translocation to the nucleus and binding of the coactivators CBP (CREB-binding protein) and p300 (18). This enables the complex to bind to target genes at the consensus sequence 5′-RCGTG-3′ (where R refers to A or G) and leads to formation of the transcription initiation complex (TIC) with subsequent expression of many genes that promote erythropoiesis, angiogenesis, glucose transport and metabolism, all needed in adaptation to low oxygen concentrations (19).

Beside the importance of hypoxia for the commensal flora, it has been shown that low oxygen conditions also impact epithelial cells by inducing secretion of several proteins into the surrounding of the cells, including cytokines and growth factors (20). Precisely, in the context of epithelial barrier function, the intestinal trefoil factors (TFFs) exhibit intestinal-specific barrier-protective features and are specifically upregulated under hypoxia through a hypoxia inducible factor HIF-1α –dependent manner (21). The molecular mechanisms of TFF function and how they achieve the barrier protection is still not fully understood. Recent publications indicate a stabilizing effect on mucosal mucins (22), induction of cellular signals that modulate cell-cell junctions of epithelia leading to increased levels of claudin-1, impairment of adherens junctions and facilitation of cell migration in wounded epithelial cell layers (23–25).

In recent years it has become appreciated that hypoxia additionally regulates the expression of an expanding but specific subset of miRNAs, termed hypoxamiRs (26, 27). miRNAs are endogenous, small non-coding RNAs that consist of 18-23 nucleotides. After transcription and subsequent maturation, the functional strand of the mature miRNA is loaded into the RNA-induced silencing complex (RISC), where it silences target mRNAs through mRNA cleavage, translational repression or deadenylation (28). miRNAs coordinate complex regulatory events relevant to a variety of fundamental cellular processes (29). Although it has been shown that miRNAs can participate in the regulation of barrier function (30), it remains unclear whether the hypoxic environment in the lumen of the gut can induce the expression of a specific subsets of hypoxamiRs which in turn will influence barrier function of the intestinal epithelium.

In the current study, we sought to investigate how hypoxia impacted the formation of a tight barrier in human intestinal epithelial cells. We found that human intestinal cells grown under hypoxic conditions more rapidly displayed barrier functions compared to cells grown under normoxia. We could correlate this improved barrier function with the faster assembly of the tight junction belt under low oxygen conditions. Through transcriptome microarray analysis, we identified three hypoxamiRs, miRNA-320a, miRNA-16-5p and miRNA-34a-5p, known to play a role in barrier formation. Using overexpression and depletion experiments, we could demonstrate that miRNA-320a acts as a key player in promoting barrier formation in human intestinal epithelial cells under hypoxic conditions. Our data demonstrates that the hypoxic condition around intestinal epithelial cells regulates the expression of a specific subsets of miRNAs which in turn participates in the establishment of a fully functional epithelial barrier. Importantly, our work highlights the importance of studying the cellular functions of intestinal epithelial cells under their physiological hypoxic environment.

## Results

### Low oxygen levels improve barrier function in human intestinal epithelial cells

The gastrointestinal tract is characterized by a steep oxygen gradient along the crypt-villus axis with high levels of oxygen at the bottom of the crypts and a low oxygen environment at the tip of the villi (10). Several studies (21, 31, 32) have shown that low oxygen concentrations can influence the barrier function of epithelial cells *in vitro* by changing gene expression profiles and inducing secretion of barrier-regulating proteins, i.e. TFFs. To investigate the mechanism by which hypoxic conditions regulate barrier function, the T84 colon adenocarcinoma-derived cell line was seeded onto transwell inserts and allowed to polarize under normoxic (21% O_2_) or hypoxic (1% O_2_) conditions. To determine the effect of hypoxia on the ability of T84 cells to form a tight barrier, transepithelial electrical resistance (TEER) measurements were performed at 24-hour intervals for five days. TEER is a well characterized method used to quickly access barrier function characterized by the rise in the electrical resistance over a cell monolayer. Similar to our previous observations (33), normoxic cells reached a polarized state and acquired a fully functional barrier function within 4-5 days post-seeding (Figure 1A). However, T84 cells cultured under hypoxic conditions established their barrier function significantly faster compared to cells under normoxic conditions, reaching a polarized state within two days post-seeding (Figure 1A). To further assess paracellular permeability and the integrity of the IEC-monolayer, the diffusion of fluorescein isothiocyanate (FITC)-labeled dextran across the epithelial monolayer was measured (Figure 1B). In this assay, when cells are non-polarized, dextran added to the apical chamber of a transwell insert is able to rapidly diffuse to the basal compartment. However, upon cellular polarization and creation of a tight barrier, the FITC-dextran is retained in the apical chamber. Results show that similar to the rapid increase in TEER measurements, T84 cells grown under hypoxic conditions are able to more quickly control FITC-dextran diffusion from the apical into the basal compartment of the transwell. This indicates that a tight barrier function has been achieved faster under hypoxia compared to normoxia (Figure 1B). This increase in barrier function was rapid and was already apparent at one day post-seeding. To determine whether the increase in the rate of polarization and barrier formation was also apparent at the level of the tight junction belt, T84 cells were seeded onto transwell inserts and the formation of tight junctions was monitored by indirect immunofluorescence of ZO-1 and by qPCR for the tight and adherens junction proteins E-Cadherin (CDH1), occludin (OCLN) and junctional adhesion molecule 1 (F11R/JAM-A). Results show that similar to the TEER and dextran diffusion assay, cells cultured under hypoxic conditions already showed, within one day of seeding, a well-defined tight junction belt characterized by the classical cobblestone pattern. On the contrary, cells grown under normoxic conditions did not have well defined tight junctions one day post seeding and this coincided with the presence of dispersed ZO-1 protein in the cytosol of the cells (Figure 1C). Additionally, mRNA expression of the junction proteins E-cadherin, occludin and JAM-A was increased under hypoxia. E-cadherin showed a higher induction initially after hypoxic culture, while occludin and JAM-A required a prolonged treatment under hypoxia to show increases in their expression (Figure 1D). All together these results suggest that hypoxia favors the establishment of barrier function in T84 cells.

**Figure 1:**
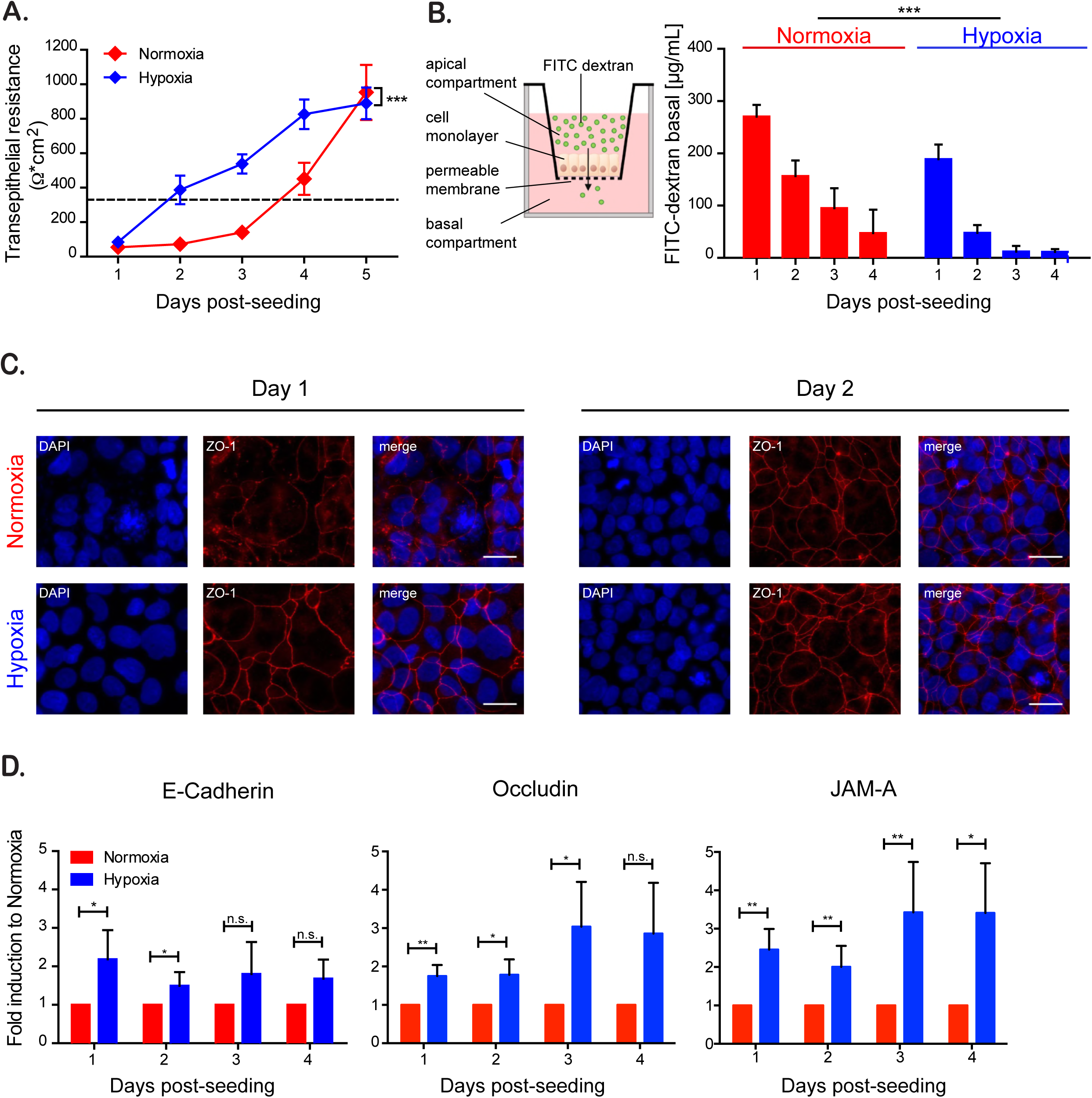
Hypoxia improves barrier function in intestinal epithelial cells. T84 cells were seeded onto transwell inserts and cultured for the indicated time under normoxic (21% O_2_) or hypoxic conditions (1% O_2_). (A) Rate of TEER increase over the cell monolayer was measured every 24 hours using the EVOM^2^ chopstick electrode. TEER greater than 330 Ohm*cm^2^ indicates complete barrier formation and is marked with a dotted line (33). (B) Paracellular permeability of the cell monolayer on transwell inserts was assessed by adding 4 kD FITC-dextran to the apical compartment (schematic overview left panel). Three hours post-incubation the basal medium was analyzed for an increase of fluorescence by spectrofluorometry (right panel). (C) T84 cultured for 24 and 48 hours under normoxic and hypoxic conditions were evaluated for the expression of the tight junction protein ZO-1 (red). Cell nuclei were stained with DAPI (blue). Scale bar indicates 25 μm. Representative image shown. (C) RNA samples of normoxic and hypoxic cultures of T84 were analyzed by qPCR for the expression of tight junction-proteins E-Cadherin, occludin and JAM-A. (A-B) Values shown represent the mean (+/-SEM) of N=9 from triplicate experiments. ***=P < 0.0001 (two-way Anova). (D) Experiments were performed in quadruplicate. Error bars indicate the standard deviation. *= P < 0.05, **= < 0.01,n.s. = not significant (one-sample t-test on log-transformed fold changes).

### Increased barrier formation induced by hypoxia is HIF-1α dependent

The main transcription factor involved in cellular response following changes in oxygenation is the hypoxia-inducible factor 1α (HIF-1α). To address whether the phenotype of faster barrier establishment under hypoxia was dependent on the activation of HIF1-α, we aimed at mimicking the hypoxic conditions using the pharmacological HIF-1α activator Dimethyloxaloylglycine (DMOG). DMOG exerts its function by inhibiting prolylhydroxylases (PHDs), which under normoxic conditions induce degradation of HIF-1α (31). Therefore, DMOG treatment of normoxic cells stabilizes HIF-1α allowing for its translocation to the nucleus and production of HIF-responsive elements (HRE) dependent gene expression (Figure 2A). To confirm that DMOG was capable of stabilizing HIF-1α in T84 cells, cells were treated with DMOG and the transcriptional upregulation of the archetypical HIF-1α -target proteins vascular endothelial growth factor (VEGF) and carbonic anhydrase 9 (CA9) were assessed by qPCR. Results show that, similar to hypoxic treatment (Supp. Figure 1A), DMOG treatment results in the significant upregulation of both VEGF and Ca9 (Suppl. Figure 1B). To determine whether HIF-1α upregulation leads to the observed increase in the rate of barrier formation, T84 cells were seeded onto transwell inserts and incubated under normoxic conditions in the presence or absence of DMOG. The barrier function was assessed by monitoring TEER in 24-hour intervals over a five-day time course. In line with our previous observations, DMOG treated cells established their barrier function faster than the solvent-treated control cells (Figure 2B). To further confirm that the observed phenotype was HIF-1α dependent, HIF-1α was knocked-down by lentiviral transduction of shRNAs (Figure 2A). Quantification of HIF-1α knockdown efficiency revealed a 75% reduction of HIF-1α mRNA compared to cells expressing a scrambled shRNA control (shScrambled) (Supp. Figure 1C). Knockdown of HIF-1α abolished the faster barrier formation under hypoxic conditions, as seen by similar TEER values under normoxic and hypoxic conditions (Figure 2C). Interestingly, cells expressing the shRNA exhibited a slower barrier formation in comparison to scrambled shRNA expressing cells even under normoxic conditions, revealing a general dependency of barrier formation on HIF-1α even in normal oxygen levels (Figure 2C). These results strongly suggest that faster establishment of barrier function in T84 cells observed under hypoxic conditions is HIF-1α dependent.

**Figure 2:**
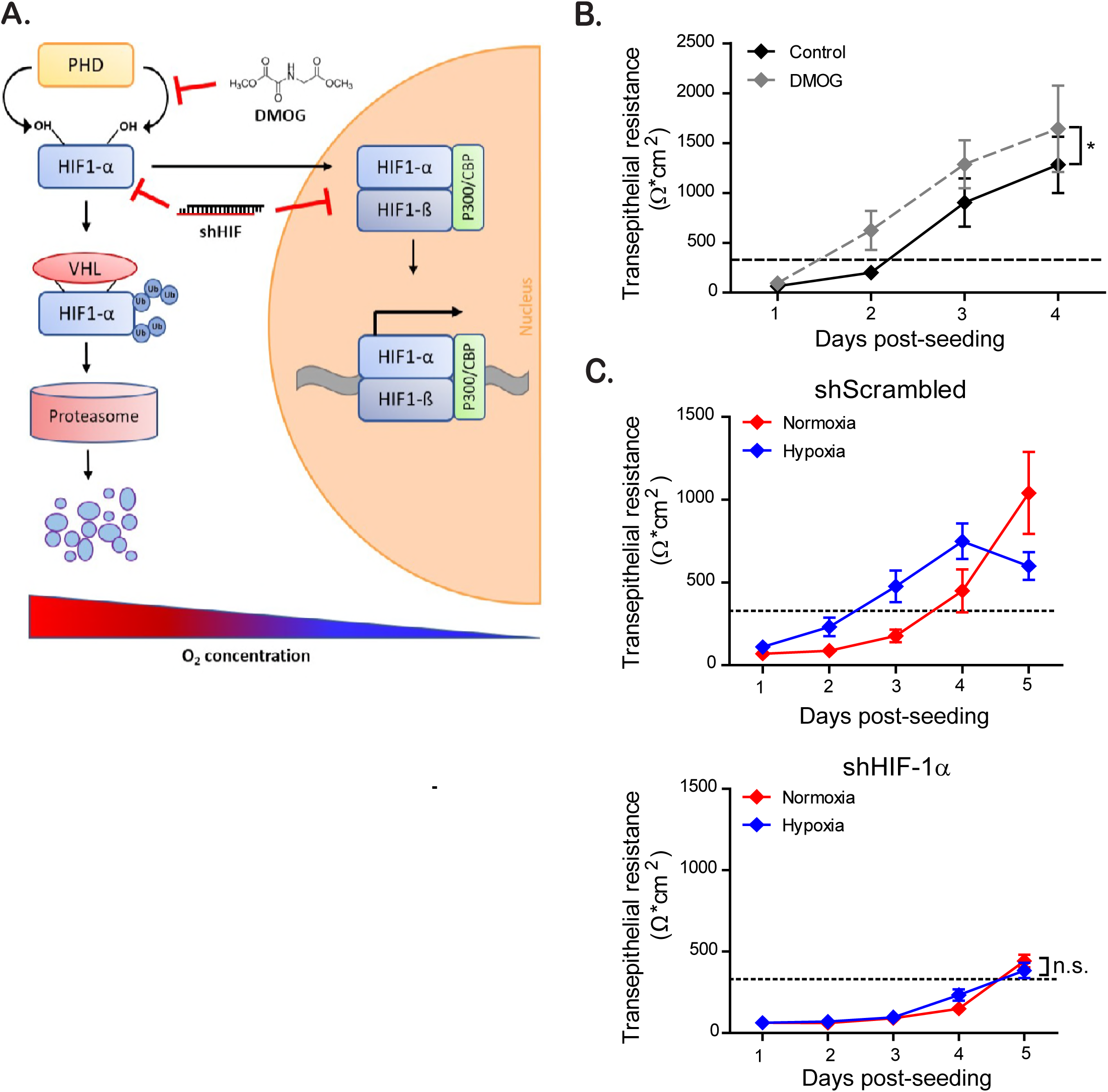
HIF-1α is responsible for faster barrier establishment under hypoxic conditions. (A) Schematic showing the regulation of the transcription factor HIF-1α at high and low oxygen concentrations. Under normoxic conditions, HIF-1α is hydroxylated at two specific proline residues by different prolyl hydroxylases (PHDs), leading to binding to the E3 ubiquitin ligase containing the von Hippel-Lindau (VHL) tumor suppressor protein. This mediates the polyubiquitination of HIF-1α and its downstream proteasomal degradation. Under hypoxic conditions, degradation is inhibited due to the lack of substrate for the PHDs, therefore stabilizing HIF-1α, leading to dimerization with its constitutively expressed β -subunit (HIF-1β) and subsequent gene expression. Pharmacological activation of HIF-1α -function by DMOG and inhibition by shRNA against HIF-1α mRNA are indicated by red arrows. (B) T84 cells were seeded on transwell inserts and incubated under normoxic conditions in the presence or absence of DMOG. TEER measurements were taken in 24-hour intervals for four days. (C) T84 cells depleted of HIF-1α through shRNA knock-down or expressing a scrambled shRNA were seeded on transwell inserts. Cells were incubated in normoxic or hypoxic conditions and TEER measurements were taken in 24-hour intervals for five days. TEER greater than 330 Ohm*cm^2^ indicates complete barrier formation and is marked with a dotted line (33). (B-C) Values shown represent the mean (+/-SEM) of N=9 from triplicate experiments. *= P:0.0417 (two-way Anova), n.s. = not significant.

### Whole transcriptome miRNA profiling reveals regulation of several miRNAs which are involved in barrier formation

Since significant differences in the barrier state between hypoxic and normoxic conditions could be observed already 24 hours after seeding, we hypothesized that the very fast changes in protein expression and barrier establishment must occur within hours after exposure of the cells to hypoxic conditions. Several proteins (21, 23) have been shown to contribute to mucosal repair and barrier formation in intestinal cells, but the role of miRNAs in finetuning gene expression involved in barrier formation has recently become appreciated (34). So far, most of these studies have only been conducted under normoxic conditions, hence overlooking the physiological hypoxic conditions of the gut. To directly address the role of miRNAs in regulating barrier functions of IECs under low oxygen conditions, miRNAome microarray analysis was employed for cells incubated under normoxic or hypoxic conditions. This allowed us to broadly screen hypoxia-regulated miRNAs, so called hypoxamiRs. By comparing the miRNA expression patterns from normoxic and hypoxic conditions we could identify a total of 108 differentially regulated hypoxamiRs of which 65 were up- and 43 were downregulated under hypoxic conditions (Figure 3A). Detailed analysis of hypoxamiRs expression revealed that upon hypoxic exposure, T84 cells highly upregulate miRNA-210-3p expression (Figure 3A). This miRNAs is a master-regulator for adaptation to low oxygen concentration (27) and is a well characterized hypoxamiRs for which expression is strongly linked to hypoxic conditions. This upregulation of miRNA-210-3p strongly suggests that T84 cells have established a hypoxia-specific transcription profile. To probe for miRNAs, which could regulate barrier function, we performed KEGG and MetaCore-driven pathway analysis allowing us to identify three potential hypoxamiRs involved in barrier function establishment (miRNA-320a, miRNA-34a-5p and miRNA-16-5p) (Figure 3B and Supp. Figure 2). miRNA-320a has been shown to be crucial for intestinal barrier integrity through modulation of the regulatory subunit PPP2R5B of phosphatase PP2A (35). Additionally, miRNA-320a was found to both target ß-catenin directly (36) and VE-cadherin through inhibition of the transcriptional repressor TWIST1 (37, 38). miRNA-34a-5p has been shown to serve as an inhibitor for the zinc-finger transcription factor Snail (39, 40), which in turn functions as a transcriptional repressor of the adherens and tight junction proteins E-Cadherin, claudins and occludin (41–43). Interestingly, we recently (44) determined that miRNA-16-5p acts as a regulator of claudin-2 expression and its expression negatively correlated with occurrence of IBS in patients, therefore playing a key role in modulating barrier function.

To validate the results of the miRNA microarray profiling, we performed qRT-PCR analysis for these specific miRNAs. As observed in our microarray approach, miRNA-210-3p, miRNA-320a, miRNA-34a-5p and miRNA-16-5p were upregulated under hypoxic conditions in T84 cells 24- or 48-hours post-seeding (Figure 4A). Since T84 cells are immortalized cells derived from carcinoma, they may show altered gene regulation, protein expression and signaling pathways. To verify that the observed hypoxia-dependent upregulation of barrier function related miRNAs was not an artefact of the cancerogenic nature of the T84 cells, stem cell-derived primary intestinal epithelial cells, so called human mini-gut organoids, were employed. Organoids are primary cell cultures and thereby retain key features like structural architecture and all major cell lineages present in the inner lining of the gut, hence mimicking the physiological organization of the human gut epithelium *in vivo* (45). In line with the results found in T84 cells, qRT-PCR confirmed upregulation of all four tested targets under hypoxic conditions 24- or 48-hours post-seeding in our human intestinal organoids (Figure 4B). Our observations made both in immortalized carcinoma derived cell lines and in primary human IECs therefore confirm the increased expression of the hypoxamiRs miRNA-320a, miRNA-34a-5p and miRNA-16-5p under hypoxic conditions in the human intestinal epithelial cells.

**Figure 3:**
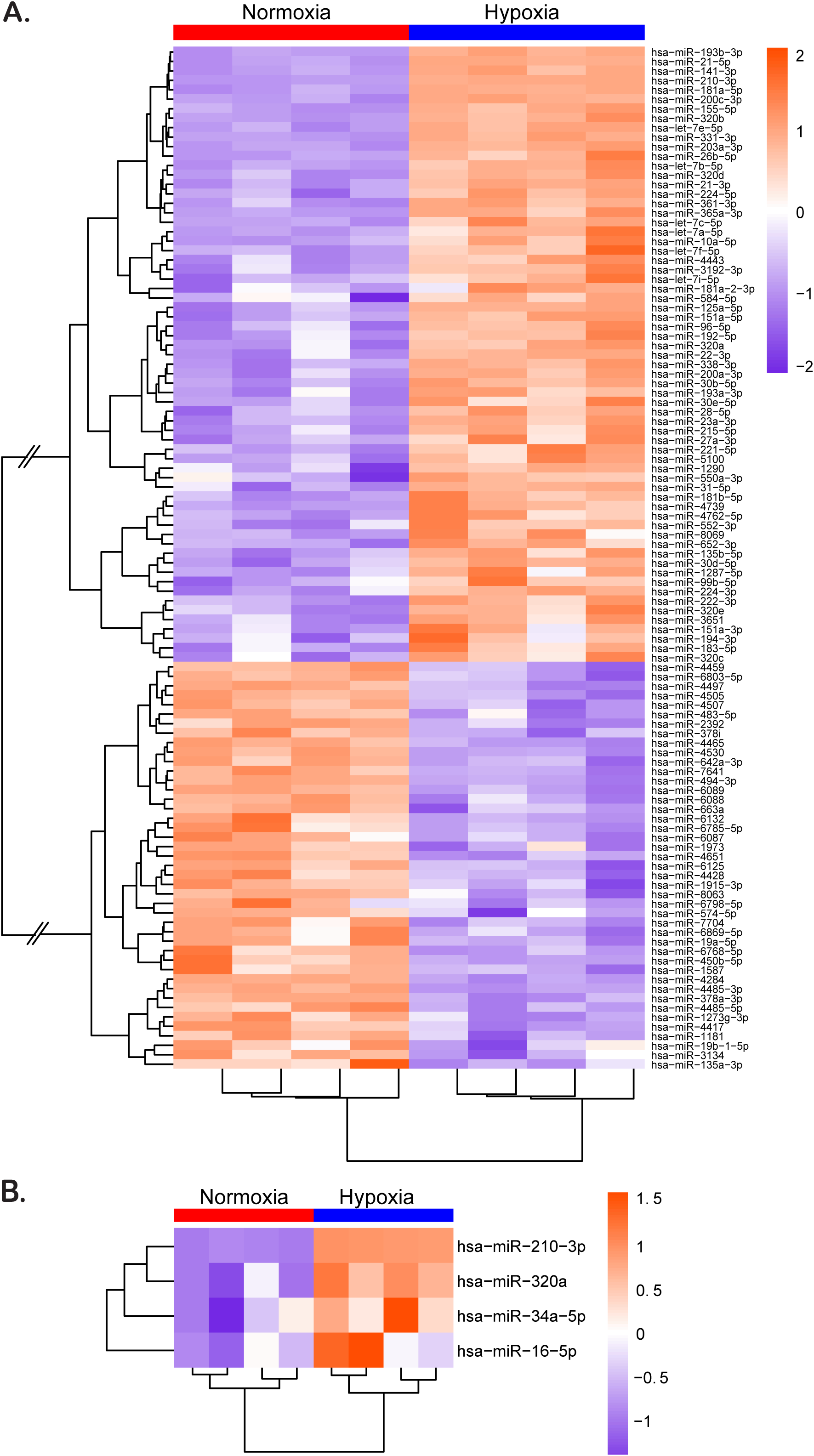
Hypoxia leads to changes in expression of several hypoxamiRs known to regulate barrier function. T84 cells were seeded on transwell inserts and incubated under hypoxic or normoxic conditions for 48 hours. miRNA was isolated and evaluated by miRNA microarray. (A-B) Heatmaps of differentially expressed miRNAs in T84 cells cultured under normoxic and hypoxic conditions. The color scale shown on the right illustrates the relative expression levels of differentially expressed miRNAs. Orange indicates up-regulated (>0), purple shows down-regulated miRNAs (<0). (A) Heatmap for 108 differentially regulated hypoxamiRs that were significantly up- or down-regulated compared to normoxic conditions. Connecting lines in the cluster dendrogram between up- and downregulated miRNAs were shortened to enable visualization (indicated by two skewed lines). (B) Heatmap of miRNA-210-3p (positive control for hypoxic conditions), miRNA-320a, miRNA-34a-5p and miRNA16-5p, identified by pathway analysis for playing a role in barrier formation. (A-B) Samples were performed in quadruplicate and the level of expression of each replicate is shown in the heatmap.

**Figure 4:**
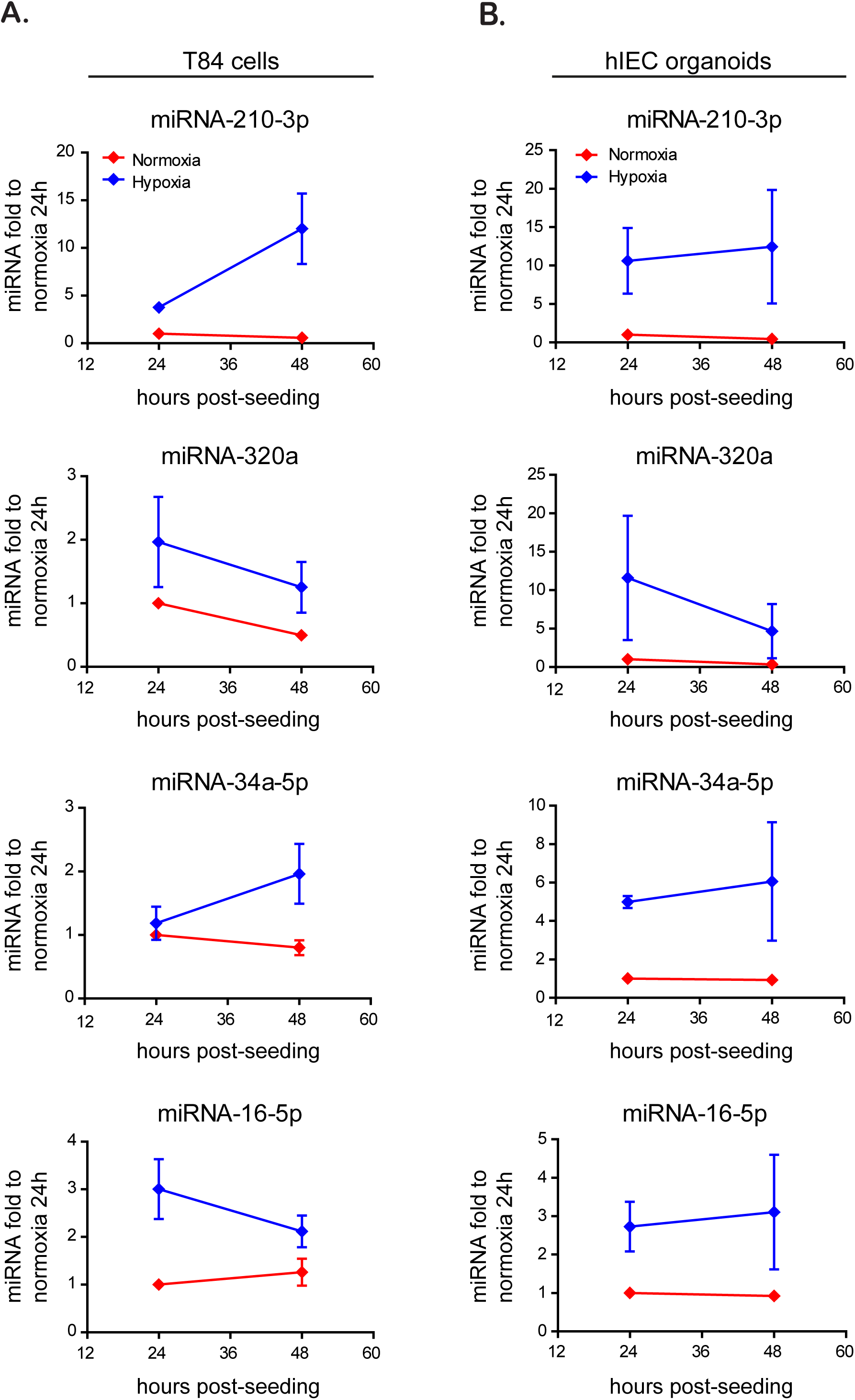
Validation of upregulated hypoxamiRs in carcinoma derived T84 cells and primary human mini-gut organoids. The expression of miRNA-210-3p (hypoxia control), miRNA-320a, miRNA-34a-5p and miRNA-16-5p was investigated 24 and 48 hours post transfer to hypoxia by qRT-PCR in (A) T84 and (B) human primary mini-gut organoids. Data was normalized to normoxic cells 24 hours post transfer. All experiments were performed in triplicate. Error bars indicate the standard error (SEM).

### Overexpression of miRNA-320a and miRNA-16-5p induces faster barrier formation in T84 cells

Our above results indicate that miRNA-320a, miRNA-34a-5p and miRNA-16-5p are upregulated under hypoxic conditions. To directly validate that these hypoxamiRs are responsible for the observed improved barrier function under hypoxia, we stably overexpressed these miRNAs in T84 cells by lentiviral transduction. Following confirmation of their overexpression using qRT-PCR (Suppl. Figure 3), miRNA overexpressing T84 cells were seeded on transwell inserts and their barrier formation was monitored by TEER measurements in 24-hour intervals (Figure 5). Results show that miRNA-320a overexpressing cells exhibited a significantly faster barrier formation in comparison to scrambled miRNA expressing cells. miRNA-16-5p over expressing cells also showed a slight but non-significant increase in barrier formation as compared to scrambled miRNA expressing cells. miRNA-34a-5p expressing cells showed no alteration in barrier formation compared to scrambled miRNA cells, even though they displayed the highest overexpression levels (Figure 5 and Suppl. Figure 3). Taken together, these data provide direct evidence for a key role of miRNA-320a in regulating barrier function in intestinal epithelial cells.

**Figure 5:**
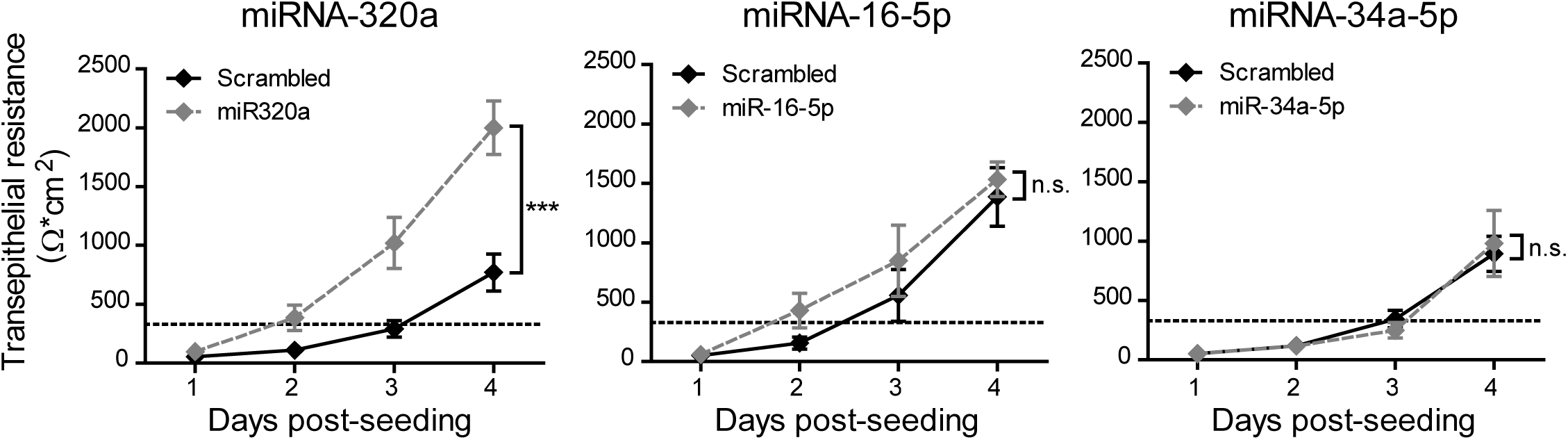
Overexpression of miRNA-320a and miRNA 16-5p induces faster barrier formation in T84 cells. T84 cells stably expressing miRNA-320a, miRNA-16-5p and miRNA-34a-5p by lentiviral transduction were seeded onto transwell inserts and barrier formation was assessed by TEER measurement in 24-hour intervals over four days. TEER greater than 330 Ohm*cm^2^ indicates complete barrier formation and is marked with a dotted line (33). Values shown represent the mean (+/-SEM) of N=6 (miRNA-16-5p & miRNA-34a-5p) or N=12 (miRNA-320a) from triplicate or quadruplicate experiments, respectively. ***= P:0.0002 (two-way Anova), n.s. = not significant.

### Inhibition of miRNA-320a expression diminishes barrier formation in T84 cells

To confirm the role of miRNA-320a in increasing barrier formation under hypoxic conditions, we generated T84 cells expressing a miRNA-320a-sponge. We confirmed through qPCR that these cells have a downregulation of miRNA-320a as the sponge binds to the miRNA and blocks its function (Suppl Figure 4). In line with our previous results, T84 cells expressing a miRNA-320a sponge displayed a slower establishment of barrier function in comparison to scrambled transduced cells under both normoxic and hypoxic conditions (Figure 6A). The effect was much more prominent under hypoxic conditions, decreasing the rate of barrier formation to the level of normoxic scrambled cells, thereby abolishing the hypoxia-dependent miRNA-320a driven barrier establishment. To confirm the role of miRNA-320a in regulating barrier function, T84 cells over expressing miRNA-320a or depleted of miRNA-320a were seeded on transwell inserts and their barrier integrity was monitored using the FITC-dextran diffusion assay. In line with our previous results, miRNA-320a overexpressing cells show a reduced flux of FITC-dextran to the basal compartment of the transwell chamber, while cells depleted of miRNA- 320a show an increased flux compared to scrambled miRNA cells (Figure 6B). Taken together, these findings strongly suggest a model where hypoxia-induced expression of miRNA-320a directly regulates the establishment of a functional barrier in the epithelial cells lining our gastrointestinal tract.

**Figure 6:**
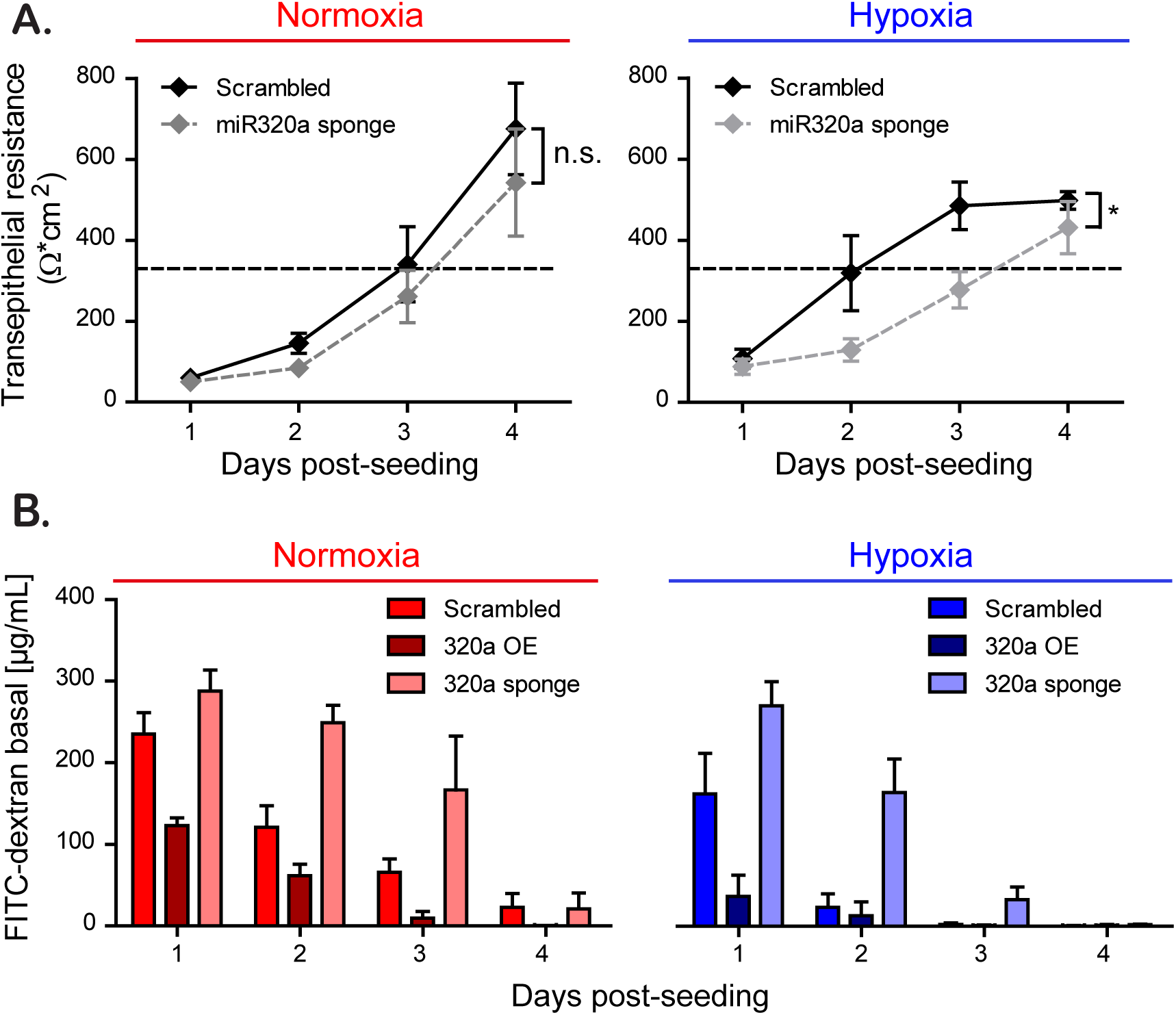
Inhibition of miRNA-320a expression diminishes barrier formation in T84 cells. (A) T84 cells stably expressing miRNA-320a sponge were seeded onto transwell inserts and barrier function was assessed by TEER measurements in 24-hours intervals over four days. TEER greater than 330 Ohm*cm^2^ indicates complete barrier formation and is marked with a dotted line (33). (B) Paracellular permeability of T84 cells overexpressing the miRNA-320a (overexpression (OE)) or the miRNA-320a sponge. Cell monolayer on transwell inserts was assessed by adding 4 kD FITC-dextran to the apical compartment and measuring fluorescence of the basal medium three hours post treatment every 24 hours for four days. Values shown represent the mean (+/-SEM) of N=12 from quadruplicate experiments (A) and N=3 from triplicate experiments (B), *= P: 0.0174 (two-way Anova).

## Discussion

In this work, we demonstrate that the physiological hypoxic environment improves intestinal epithelial barrier function of T84 cells as shown by the faster establishment of transepithelial electrical resistance, by the more rapid decrease in barrier permeability to FITC-dextran, as well as by the faster establishment of the tight junction belt compared to normoxic conditions. Using pharmacological inhibitor and knock-down approaches, we could show that this increased barrier function is dependent on the hypoxia regulator HIF-1α Additionally, using a miRNA microarray approach we identified miRNA-320a as a key miRNA induced under hypoxia being directly responsible for regulating barrier functions in human intestinal epithelial cells. We could demonstrate that its overexpression is sufficient to promote barrier function in epithelial cells while interfering with its expression under hypoxic conditions counteracts the hypoxia-mediated barrier formation establishment. Together our results show that miRNA-320a is a hypoxia-induced miRNA which plays a key role in regulating barrier function in human intestinal epithelial cells.

The importance of hypoxic conditions in regulating barrier function in intestinal epithelial cells has been previously studied and several potential mechanisms highlight the central role of the transcription factor HIF. It was shown that specific shRNA-mediated knock- down of HIF-1β in T84 and Caco-2 cells resulted in the decrease of claudin-1 expression on mRNA and protein level accompanied by defects in barrier function and abnormal morphology of tight junctions (46). This is thought to be a direct effect from the HIFs themselves as HIF responsive elements have been identified in the promoter region of claudin-1 (46).

One of the best characterized means by which hypoxia induces barrier formation involves the HIF-dependent expression of the intestinal trefoil factor (TFFs). The trefoil factor family consists of three peptides: TFF1, TFF2 and TFF3; all three are widely distributed in the gastrointestinal tract and are present in virtually all mucosal membranes (47). Recently, TFFs have been shown to induce a stabilizing effect on mucosal mucins (22). Additionally, the Van- Gogh-like protein 1 (Vangl1) was identified as a downstream effector of TFF3 and described to mediate wound healing in IECs, thereby promoting recovery of barrier function under condition of local loss of epithelium integrity (23). Importantly, TFF3 also regulates the expression of tight junctions and adherens junctions in IECs by elevating the levels of claudin-1 and downregulating the expression of E-cadherin (24). This further activates the phosphatidylinositol 3-kinase (PI3K)-Akt signaling pathway, which leads to an increase in barrier function and altered proliferation of cells in the intestinal epithelium (25, 48). Extensions of these studies *in vivo* revealed the protective role of TFFs on intestinal permeability and barrier function, as both administration of TFFs as well as administration of a novel prolyl hydroxylase (PHD) inhibitor (FG-4497) were protective and had a beneficial influence on clinical symptoms (weight loss, colon length, tissue TNFα) in a mouse colitis model (49, 50). Correspondingly, HIF-1α was found to be highly expressed in Crohn’s Disease and ulcerative colitis patients (51) and seems to play a protective role in inflammatory bowel disorders through improvement of epithelial barrier function (52). It has been suggested that HIF-1α helps to control intestinal inflammation by interacting with the inflammation transcription factor nuclear factor-kappa B (NF-?B) (53).

To date, most of the work aimed at understanding the effect of hypoxia on barrier function in the gut has focused on the transcripts and proteins that are induced under hypoxia. In the emerging field of miRNA, several miRNAs have been identified as potential regulators of barrier function. However, to the best of our knowledge, these miRNAs were not studied under hypoxic conditions but in normal cell culture conditions or in patient samples with inflammatory diseases. For example, McKenna et *al*. demonstrated that claudin-4 and claudin-7 were not expressed in the apical membrane of intestinal epithelial cells in Dicer 1-deficient mice, resulting in impaired intestinal barrier function thus strongly supporting the importance of miRNA regulation in barrier formation (54). Additionally, overexpression of miRNAs has been linked to a regulation of barrier function in intestinal epithelial cells (35, 55). miRNA-31 was found to increase the TEER by decreasing the transepithelial permeability through interaction with tumor necrosis factor superfamily member 15 (TNFSF15) in Caco2-BBE cells (56). Of note, TNFSF15 is a well-known risk gene involved in the pathogenesis of irritable bowel syndrome (IBS) and inflammatory bowel disease (57, 58). hsa-miRNA-26b was found to regulate the Ste20-like proline/alanine rich kinase (SPAK) involved in epithelial barrier integrity (59) and overexpression of miRNA-21 in patients with ulcerative colitis has been associated with the impaired intestinal epithelial barrier function through targeting the Rho GTPase RhoB (60). We recently identified miRNA-16 and miRNA-125b, as being downregulated in patients suffering from IBS with diarrhea and determined that these two miRNAs modulated the tight junction proteins claudin-2 and cingulin (44).

Similar to our work, miRNA-320a was previously reported to play a role in barrier function under normoxic conditions. Cordes et *al*. could show a functional role of miRNA-320a in stabilizing the intestinal barrier function through reinforcement of barrier integrity in T84 cells and in a murine colitis model (35). They suggest that this is due to a potential modulation of the tight junction complex during intestinal inflammation. However, they did not address how different oxygen concentration could influence expression of this hypoxamiR. Our miRNA expression profiling showed an upregulation in all members of the miRNA-320 family under hypoxic conditions. We further demonstrate that, as a result of the induced expression of miRNA-320, hypoxic conditions favor barrier function of intestinal epithelial cells. As such we propose that the hypoxic environment present in the lumen of the gut impacts barrier functions not only via direct HIF-mediated regulation of tight junction and adherens proteins expression but also through a miRNA-based regulation of cell-cell contact formation.

To conclude, our work further emphasizes the importance of studying intestinal epithelial cells in their physiological environment. On the one hand, hypoxia directly influences the cell biology of the mucosal layer by regulating cell to cell contact, migration, stem-cellness and metabolism. On the other hand, a low oxygen concentration is critical for the establishment and maintenance of a stable microbiota. As such, given the growing interests in understanding both host/commensal interactions in health and diseases and the complex interplay between host and pathogens in the gastrointestinal tract, it is critical to integrate the impact of local oxygen concentration and fluctuation in regulating/altering these molecular processes.

## Acknowledgements

This work was supported by a research grant from Chica and Heinz Schaller Foundation and Deutsche Forschungsgemeinschaft (DFG) in SFB1129 (Project 14) to SB. This project has received funding from the European Union’s Seventh Framework Programme under grant agreement no 334336 (FP7-PEOPLE-2012-CIG). MS was supported by the Brigitte-Schlieben Lange Program from the state of Baden Württemberg, Germany and the Dual Career Support from CellNetworks, Heidelberg, Germany. We would like to thank the lab of Hanno Glimm, NCT, Heidelberg for intestinal tissue samples, Himanshu Soni and Björn Tews for providing the HIF-1a shRNA lentivirus construct and the Genomics and Proteomics core facility of the German Cancer Research Center for their preparation and processing of the miRNA microarray samples.

## Material & Methods

### Cell Lines

T84 human colonic adenocarcinoma cells (ATCC CCL-248) were cultured in GibCo’s Dulbecco’s Modified Eagle Medium/F-12 Nutrient Mixture (1:1), supplemented with 10 % fetal bovine serum (FBS), 100 U/mL penicillin and 100 μg/mL streptomycin (GibCo) in collagen coated T25 cell culture flasks. The cells were kept in a constant humid atmosphere containing 37°C, 5% CO_2_ and either 21% oxygen (normoxia) or 1% oxygen (hypoxia). HEK293T human embryonic kidney cells (ATCC CRL 3216) and cultured in Iscove’s modified Dulbecco’s medium supplemented with 10% FBS and 100 U/mL penicillin and 100 μg/mL streptomycin. Cells were grown at 37°C in a humidified atmosphere containing 5% CO_2_. Human intestinal epithelial organoids were isolated from biopsy tissue provided by the University Hospital Heidelberg as described before (61). This study was carried out in accordance with the recommendations of the University Hospital Heidelberg with written informed consent from all subjects in accordance with the Declaration of Helsinki. All samples were received and maintained in an anonymized manner. The protocol was approved by the “Ethics commission of the University Hospital Heidelberg” under the protocol S-443/2017. In short, resected intestinal tissue was incubated with 2 mM EDTA in PBS for 1 hour at 4°C. Intestinal crypts containing the Lgr5+ stem cell niche were isolated after 2 mM EDTA treatment, washed with ice cold PBS and resuspended in Matrigel. The Matrigel was then overlaid with basal medium (Advanced DMEM/F12, supplemented with 1% penicillin/streptomycin, 10?mM HEPES, 50% v/v L-WRN conditioned media (ATCC #CRL-3276, expressing Wnt3A, R-spondin and Noggin), 1x B-27 (Life technology), 1x N-2 (Life technology), 2 mM GlutaMax (Gibco), 50?ng/mL EGF (Invitrogen), 1?mM *N*-acetyl-cysteine (Sigma), 10?mM nicotinamide (Sigma), 10 μM SB202190 (Tocris Bioscience) and 500?nM A-83-01 (Tocris)) and cultured at 37°C, 5% CO_2_ and 21% or 1% oxygen.

### Antibodies/Reagents

Mouse monoclonal antibody against ZO-1 (Invitrogen #339100) was used at a 1/100 dilution for immunostaining. Secondary antibodies were conjugated with AF568 (Molecular Probes) and directed against the animal source. ProLong Gold Antifade containing DAPI was obtained from Thermo Fisher Scientific. 4 kDa FITC-labelled dextran and Dimethyloxalylglycine (DMOG) was obtained from Sigma-Aldrich.

### Monitoring Transepithelial Electrical Resistance

To monitor barrier function, 1 × 10^5^ T84 cells were grown on transwell filters (6.5 mm polycarbonate membrane, 3 μm pore size; Corning). The medium was changed one day post seeding and subsequently every second day. Transepithelial resistance was measured with the EVOM^2^ chopstick electrode. T84 cells were considered to have a completely formed barrier when being also fully polarized. Full polarization in our setting was reached with a TEER of 1000 Ω (33). Taking into account the surface of the membrane, reaching a value of 330 Ω*cm^2^ indicated full barrier function (62).

### Fluorescent flux assay using fluorescein isothiocyanate (FITC)-labeled dextran

1×10^5^ T84 cells were grown on collagen coated transwell filters under normoxic and hypoxic conditions. Every 24 hours, 2 mg/mL FITC-labelled dextran was added to the apical compartment and media was collected from the basal compartment three hours post-treatment. Increase of fluorescence in the basal media was measured with the FLUOstar Omega spectrofluorometer (BMG Labtech) at an excitation wavelength of 495 nm and an emission wavelength of 518 nm. As a positive control, the fluorescence of a 100 μL aliquot of a collagen coated but cell-free transwell filter was measured to assess maximum diffusion of FITC-labeled dextran.

### Immunofluorescence staining

1×10^5^ T84 cells were grown on transwell filters. At the indicated times post-seeding, the polycarbonate membrane was removed from the transwell holder, rinsed once in PBS and fixed in 2% PFA for 20 min. PFA was removed, cells were washed 3x with PBS and permeabilized with 0.5% Triton X-100 (v/v) at RT for 15 min. After blocking with 3% BSA-PBS for 1?hour at RT, cells were incubated with primary antibody against ZO-1 in 3% BSA-PBS for 1?hour at RT. Cells were then washed with 0.1% Tween-20-PBS (v/v) followed by incubation with the secondary goat anti-mouse Alexa 568 antibody diluted in 1% BSA at RT for 45 min. After 45 min, cells were subjected to 3x washing with 0.1% Tween-20-PBS. The membrane was then briefly rinsed in Millipore H_2_ O and mounted onto glass slides using ProLong Gold Antifade reagent with DAPI. Samples were imaged on a Nikon Eclipse Ti-S inverted microscope using a 40x oil objective.

### RNA Isolation, cDNA, and qPCR

Total RNA was purified from lysed T84 colonic adenocarcinoma cells or intestinal organoids using the NucleoSpin RNA extraction kit by Marchery-Nagel following the manufacturer’s instruction. 100-250 ng total RNA was reversed transcribed into cDNA using the iScript cDNA Synthesis kit as per manufacturer’s instruction (BioRad Laboratories). qRT-PCR was performed using the Bio-Rad CFX96 Real-Time PCR Detection System and SsoAdvanced Universal SYBR Green Supermix (Bio-Rad). The data was analyzed with the Bio-Rad CFX Manager 3.0, using the housekeeping gene HPRT1 for normalization. Expression of E-Cadherin, occludin, Jam-A, VEGF and CA9 were analyzed using specific primers for the respective human sequence. The expression levels of the investigated genes were calculated as ΔΔCq, normalizing to normoxic control samples and to the normalizing genes.

**Table 1:**
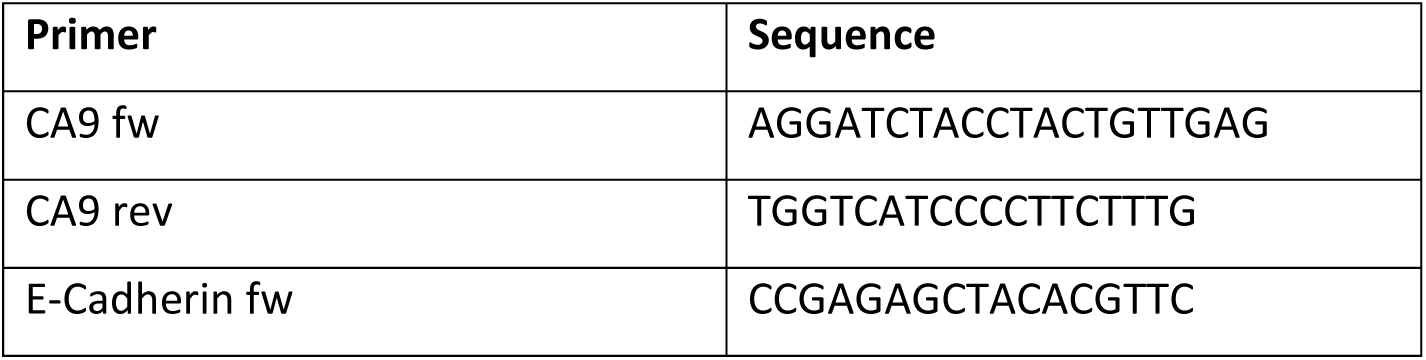
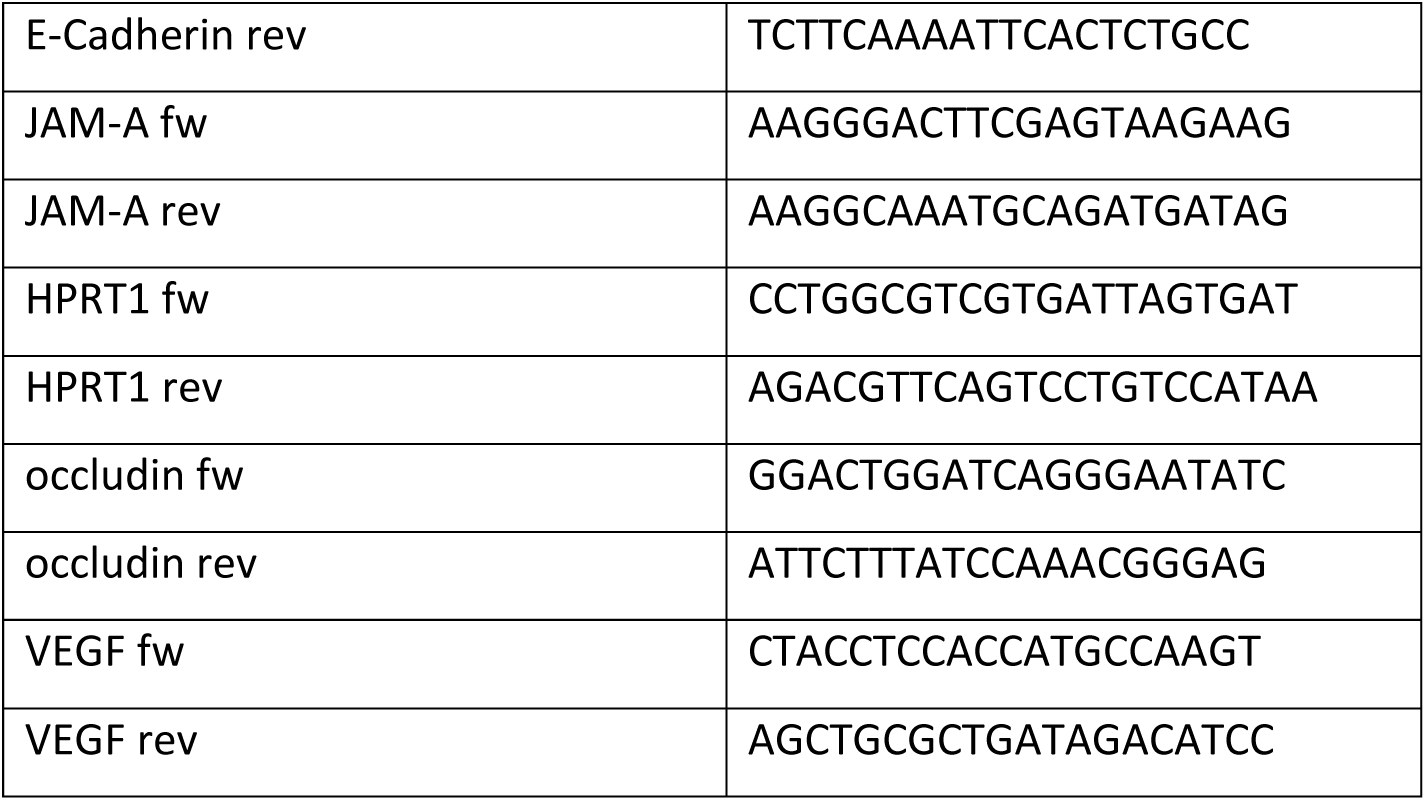
Primer Sequences qRT-PCR

### miRNA microarray

Expression of miRNA’s under normoxic and hypoxic conditions was analyzed by extracting total RNA including miRNA using the miRNeasy Mini Kit by Qiagen according to the manufacturer’s instructions. Microarray analysis was performed using the Agilent human miRNA v21 microarray chip. Quantile normalized miRNA expression values were log2-transformed and differentially expressed miRNAs between experimental conditions were identified using the empirical Bayes approach based on moderated t-statistics as implemented in the Bioconductor package limma. P-values were adjusted for multiple testing using the Benjamini-Hochberg correction to control the false discovery rate. Adjusted p-values below 5% were considered statistically significant. For heatmap display, miRNAs were scaled across samples, and hierarchical clustering of samples and miRNAs was performed using euclidean distance and Ward’s linkage. Analyses were carried out using R 3.348, with add-on package pheatmap. Target genes of significantly regulated miRNAs were retrieved from miRTarBase database v6.1 using Bioconductor package multiMiR (63). Overrepresentation of KEGG pathways was tested with limma functions kegga and goana. P-values were adjusted for multiple testing using the Benjamini-Hochberg correction. Subsequent pathway analysis was performed using the MetaCore™ software.

### miRNA validation

For further validation of miRNA-210-3p, miRNA-320a, miRNA-34a-5p and miRNA-16-5p, total RNA including miRNA was transcribed into cDNA using the miScript II RT Kit (Qiagen). After cDNA-synthesis, qRT-PCR was performed using the miScript SYBR^®^ Green PCR Kit (Qiagen) and the respective miScript Primer Assays (Qiagen) on the Bio-Rad CFX96 Real-Time PCR Detection System, normalizing to RNU6-2 as a housekeeping snRNA. The fold of expression of the investigated miRNAs were calculated as ΔΔCq, normalizing to normoxic control samples and to the housekeeping snRNA.

### Production of lentiviral constructs expressing miRNAs and shRNA against HIF-1α

Oligonucleotides encoding the sequence for mature miRNA-16-5p, miRNA-34a-5p, and miRNA-320a were designed according to protocol “Lentiviral Overexpression of miRNAs” (64), oligonucleotides encoding the sequence for HIF-1α knockdown were designed from the TRC library, cloneID: TRCN0000003808 (Table 2). Annealed oligonucleotides were ligated with the AgeI-HF and EcoRI-HF digested pLKO.1 puro vector (Addgene #8453) using the T4 DNA Ligase (New England Biolabs) and the resulting plasmids were transformed into *E. coli* DH5α-competent cells. Amplified plasmid DNA was purified using the NucleoBondR PC 100 kit by Marchery-Nagel following the manufacturer’s instruction.

**Table 2:**
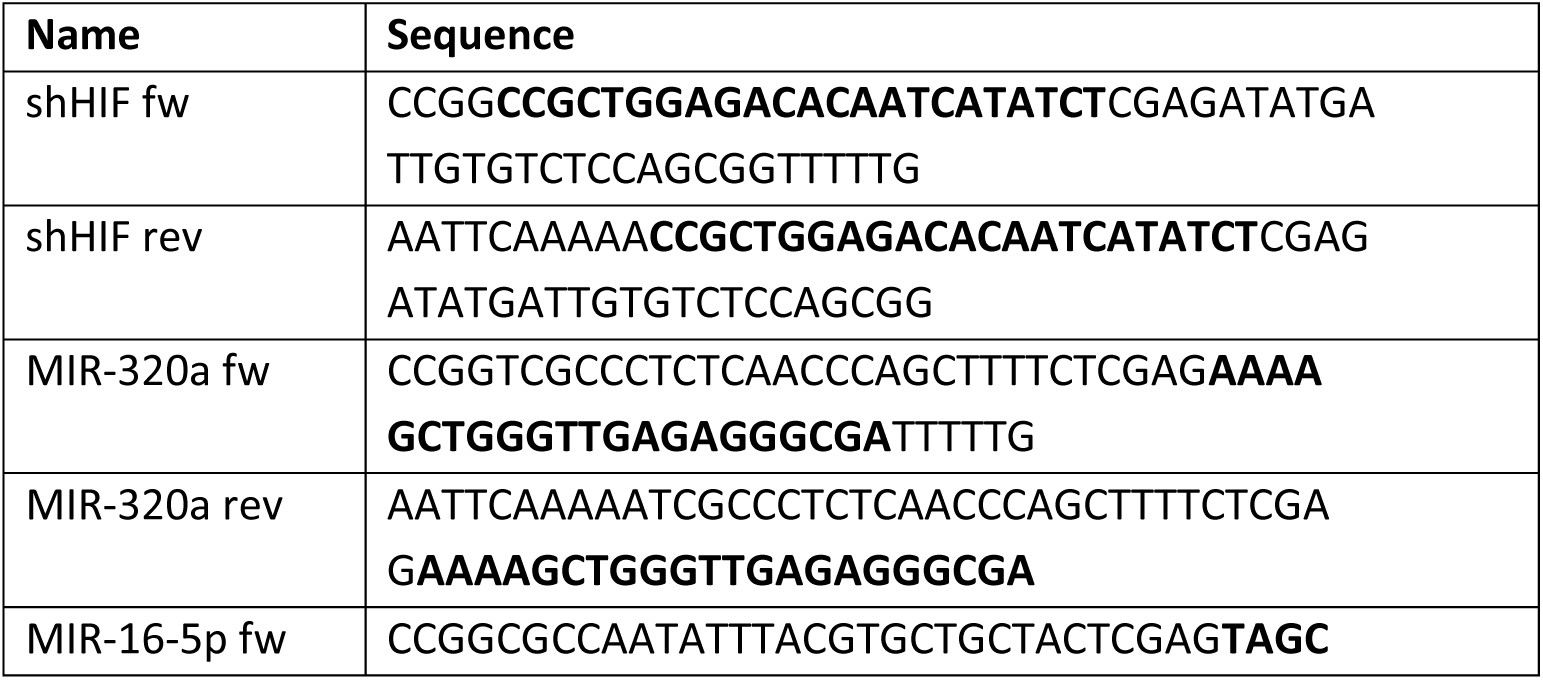
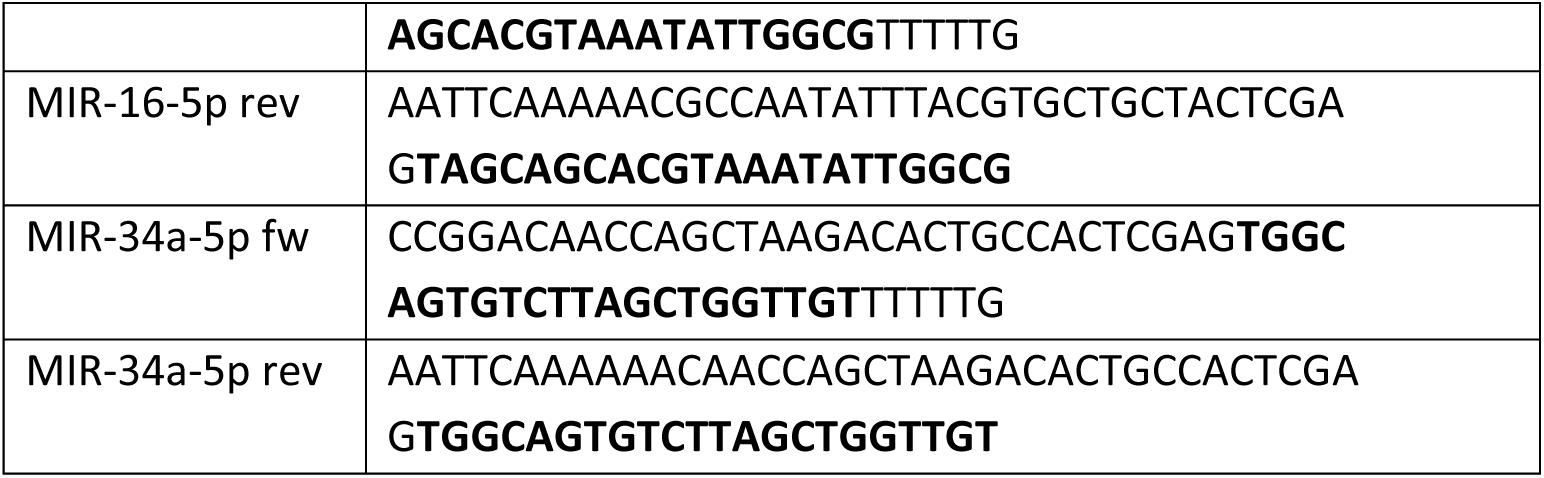
Oligonucleotides for shRNA and miRNA expression. Bold characters mark the respective target or miRNA sequence.

### Lentivirus production and selection of stable cell lines

HEK293T cells were seeded on 10 cm^2^ dishes and allowed to adhere for 2 days. When cells reached 70-80% confluence they were transfected with 4 μg of pMD2.G (Addgene #12259), 4 μg of psPAX2 (Addgene #12260) and 8 μg of purified pLKO.1 plasmid containing the shRNA or miRNA constructs. Cell supernatant containing generated lentivirus was harvested 48-72 h post-transfection, filtered through a 45 μM Millex HA-filter (Merck Millipore) and purified by ultracentrifugation at 27,000x g for three h. For lentiviral transduction, 3×10^5^ T84 cells were seeded onto collagen coated 6-well plates. After 24 h, medium was replaced with 4 mL medium containing 20 μL of the purified lentivirus or lentivirus encoding the 320a-sponge (MISSION^®^ Lenti microRNA Inhibitor, Human, Sigma, #HLTUD0470). Two to three days after transduction, medium was supplemented with 10 μg/mL puromycin for selection of successfully transduced cells.

## Supplementary Information

**Supplementary Figure 1.**
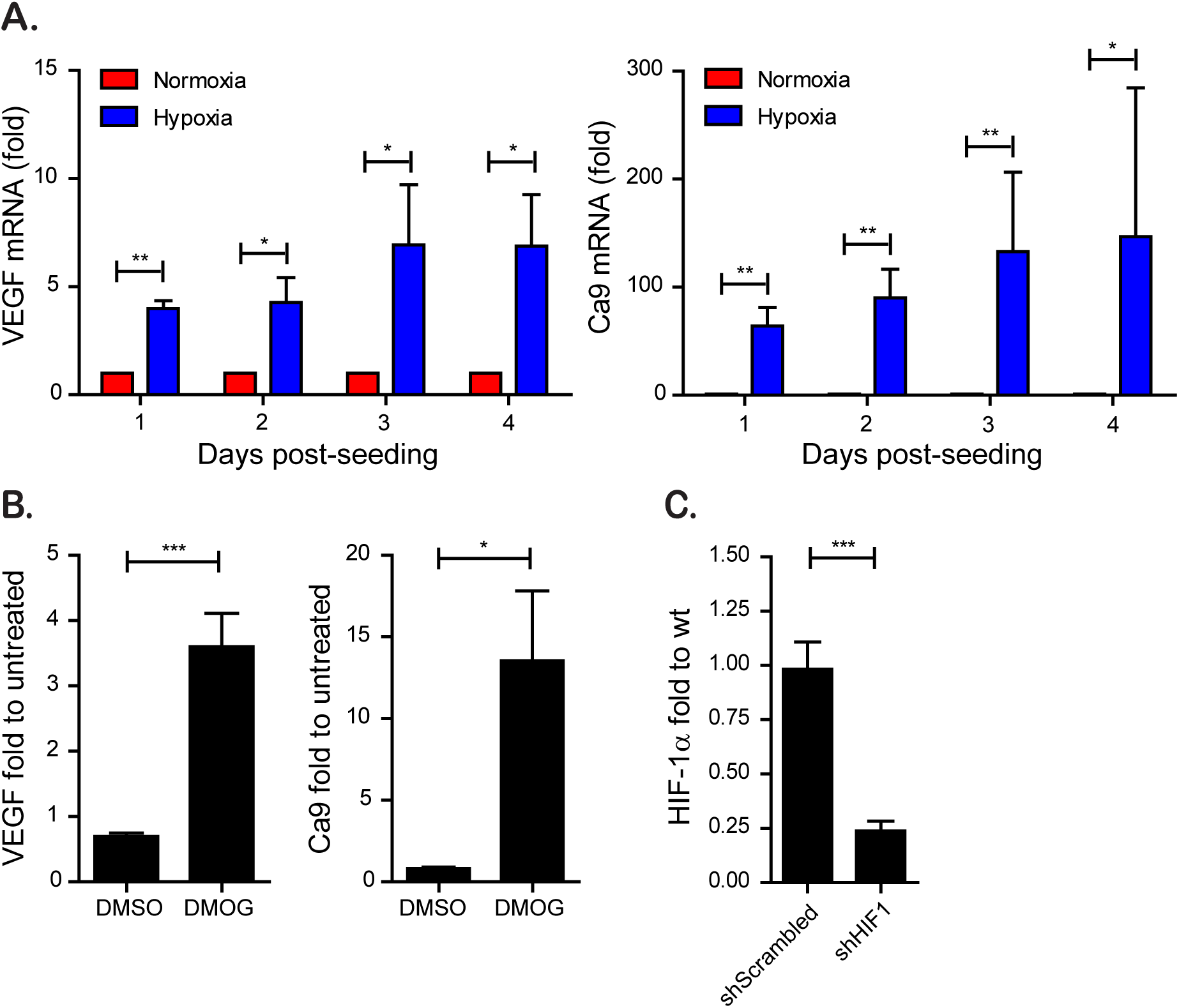
HIF-1α modulation through pharmacological treatment and shRNA knock-down. (A) RNA samples of normoxic and hypoxic cultures of T84 taken in 24-hour intervals for four days were analyzed by qPCR for the expression of the hypoxia-induced genes VEGF and Ca9. (B) T84 cells were seeded on transwell inserts and incubated under normoxic conditions in the presence or absence of DMOG. RNA was isolated and the upregulation of VEGF and Ca9 were evaluated by qPCR. (C) T84 cells expressing a shRNA against HIF-1α were evaluated for their expression of HIF-1α. (A-C) Values shown represent the mean plus standard deviation of three (A) or four (B,C) independent experiments, *= P < 0.05, **= P < 0.01, ***= P < 0.001, n.s. = not significant (one-sample t-test on log-transformed fold changes).

**Supplementary Figure 2.**
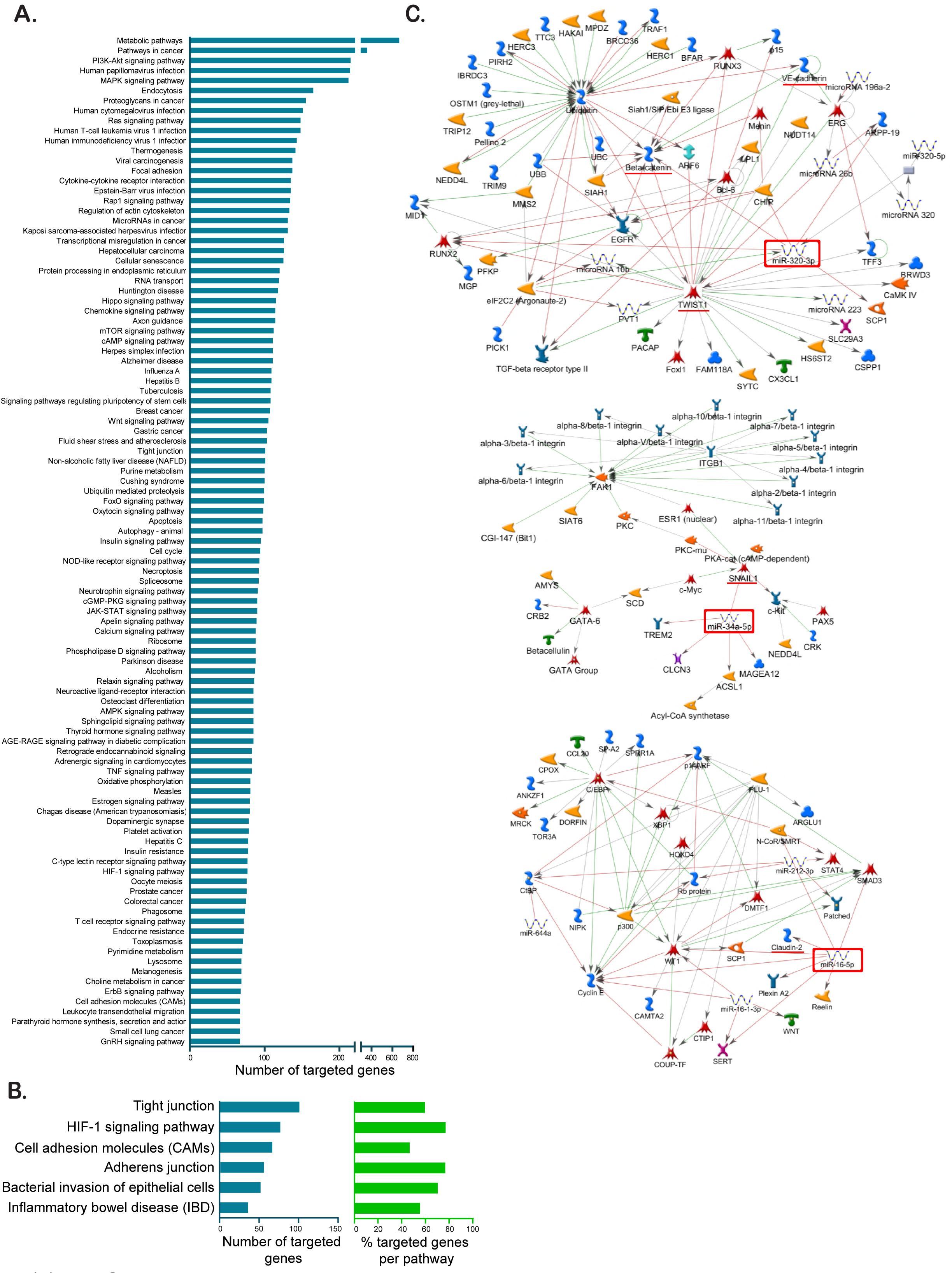
Pathway analysis by KEGG and MetaCore reveals miRNA-320a, miRNA-34a-5p and miRNA-16-5p as regulators of tight- and adherens junction proteins. (A) Target genes of significantly regulated miRNAs were retrieved from miRTarBase database v6.1 and subjected to KEGG pathway analysis. The 100 most targeted pathways by number of targeted genes are shown.(B) Number of targeted genes and percentage of targeted genes per pathway for barrier function related pathways. (C) MetaCore-driven pathway analysis identified three potential hypoxamiRs involved in barrier function establishment. Interaction maps are shown for (A) miRNA-320a, (B) miRNA-34a-5p and (C) miRNA-16-5p. miRNA of interest is marked by a red square, targeted proteins involved in barrier formation are underlined in red.

**Supplementary Figure 3.**
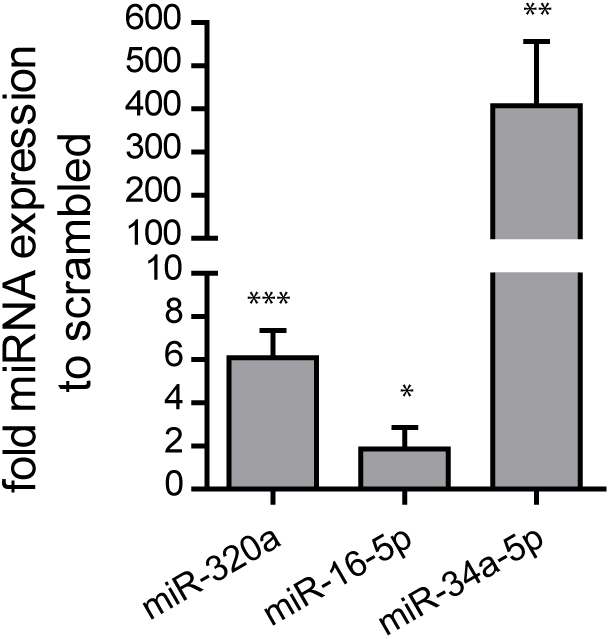
T84 cells overexpress miRNAs after lentiviral transduction. T84 cells were selected to overexpress miR-320a, miR-16-5p, and miR-34a-5p through lentivirus transduction. Cells were harvested and the overexpression of each miRNA was evaluated by miScript PCR. Values shown represent the mean plus standard deviation of three independent experiments. *= P < 0.05, **= P < 0.01, ***= P < 0.001 (one-sample t-test on log-transformed fold changes).

**Supplementary Figure 4.**
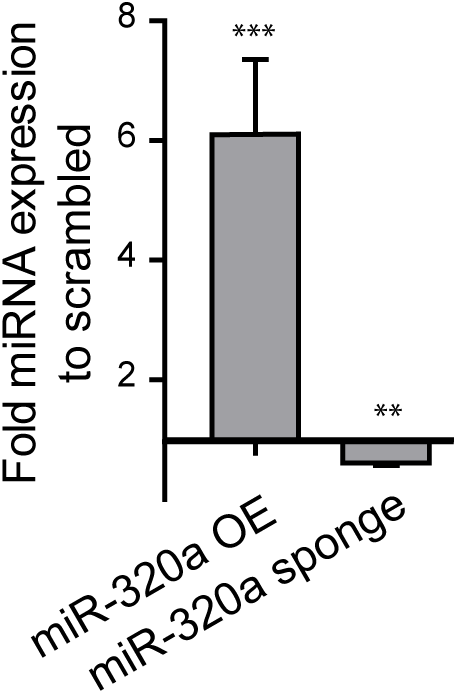
Confirmation of miRNA-320a and miRNA-320a sponge expression in T84 cells. T84 cells overexpressing (OE) miRNA-320a or depleted of miRNA-320a by expression of a sponge were evaluated by miScript PCR. Values shown represent the mean plus standard deviation of three independent experiments. *= P < 0.05, **= P < 0.01, ***= P < 0.001 (one-sample t-test on log-transformed fold changes).

